# Topographic functional interactions between caudate and cortex are organized by network and degrade with age

**DOI:** 10.1101/2019.12.23.887398

**Authors:** Jonathan F. O’Rawe, Hoi-Chung Leung

**Affiliations:** Integrative Neuroscience Program, Department of Psychology, Stony Brook University

**Keywords:** cortico-striatal connectivity, functional organization, topography, aging, flexible behavior

## Abstract

The striatum is postulated to play a role in gating cortical processing during goal-oriented behavior. However, the underlying circuit structure for striatal gating remains unclear. Deviating from previous approaches which typically treat the striatum as a homogenous structure or small compartments, we took a functional connectivity approach that utilizes the entire anatomical space of the caudate nucleus and examined its functional relationship with the cortex and how that relationship changes with age. We defined the topography of the caudate functional connectivity with the rest of the brain using three publicly available resting-state fMRI data samples. There were several key findings. First, our results revealed two stable gradients of connectivity patterns across the caudate: medial-lateral (M-L) and anterior-posterior (A-P) axes, which supports findings in previous anatomical studies of non-human primates that there is more than one organizational principle. Second, the differential connectivity patterns along the caudate’s M-L gradient were not limited to single structures but rather organized with respect to large-scale neural networks; in particular, networks associated with internal orienting behavior are closely linked to the medial extent of the caudate whereas networks associated with external orienting behavior are closely linked to the lateral extent of the caudate. Third, we found a decrease in the integrity of M-L organization with healthy aging which was associated with age-related changes in behavioral measures of flexible control. In sum, the caudate shows a topographic organization with respect to large-scale networks in the human brain and changes this organization seem to have implications for age-related decline in flexible control of behavior.

## Introduction

Clinical and basic sciences have found a critical role of the striatum in gating, a unique function of selecting action plans to optimize goal attainment (Grahn, Parkinson, & Owen, 2008; Mink, 1996). This is evident in movement disorders such as Parkinson’s Disease and Huntington’s Disease, both of which express symptoms stemming from dysfunctional motor gating and have pathology of the striatum (Bernheimer, Birkmayer, Hornykiewicz, Jellinger, & Seitelberger, 1973). Individuals with these neurological diseases also demonstrate cognitive symptoms, suggesting a much more complex outcome of striatal dysfunction (Brandt, Folstein, & Folstein, 1988; Cooper, Sagar, Jordan, Harvey, & Sullivan, 1991; Janvin, Larsen, Aarsland, & Hugdahl, 2006; Lawrence et al., 1998).

Previous studies have consistently shown that striatal functioning is closely associated with updating and switching demands during complex behavioral tasks (Badre & Frank, 2012; Chatham, Frank, & Badre, 2014; Frank, Loughry, & O’Reilly, 2001; O’Reilly, 2006). In particular, empirical evidence from lesions in the rodent striatum (Yin, Knowlton, & Balleine, 2005, 2006), and neuroimaging experiments in humans demonstrated a clear selective role of the caudate nucleus in tasks dependent on goal oriented learning (e.g., action-outcome association rather than stimulus-response associations) (Chiu, Jiang, & Egner, 2017; Haruno & Kawato, 2006). Indeed, cognitive syndromes in both Huntington’s and Parkinson’s disease have been more specifically related to caudate dysfunction (Berent et al., 1988; Ekman et al., 2012; Lewis, Dove, Robbins, Barker, & Owen, 2003).

Understanding the organization of the caudate nucleus is a critical step towards understanding the computations necessary to gate goal directed behavior, as anatomical and functional organization provides constraints the computations that can be performed (Jbabdi, Sotiropoulos, & Behrens, 2013; Thivierge & Marcus, 2007). Systematic neuroanatomical tracing studies suggested that the connections between the neocortex and the caudate are extensive and systematically organized. Early anatomical findings using silver impregnation of degenerating axons following systematic lesioning of the cortex were fairly consistent in their results: finding that cortical projections to the caudate were fairly restricted along the rostral-caudal axis, such that more caudal cortex projects to more caudal caudate, and vice versa for the rostral projections (Carman, Cowan, Powell, & Webster, 1965; Kemp & Powell, 1970; Webster, 1965). However, anterograde tracing from the cortex demonstrated far less restriction along the rostral-caudal axis, with several studies have found a consistent organization across the medial-lateral and the dorsal-ventral axes (Ferry, Öngür, An, & Price, 2000; Selemon & Goldman-Rakic, 1985). It was further shown that that areas which share corticocortical connections also show colocalization in projections to the caudate (Parthasarathy, Schall, & Graybiel, 1992; Selemon & Goldman-Rakic, 1985; Yeterian & Van Hoesen, 1978). Accordingly, it may be the case that caudate organization can be understood more comprehensively on the basis of a functional network architecture, with discrete nodes of a network compressing information through the striatothalamic pathways for integration with other cortical nodes of the same network. With such functional organization, adaptive behavior may arise quite efficiently via systematic dampening or enhancing of cortical response along different nodes of a network, by virtue of reward contingency (Bar-Gad, Morris, & Bergman, 2003).

A consistent topographic relationship of frontal cortex projections into the striatum has been shown previously in animal models (Haber, 2003). This topographic relationship tracks along a diagonal path from dorsolateral to ventromedial striatum, in which more pure motor processes, such as those from precentral sulcus, project to dorsolateral areas of the striatum, while higher order control and reward processes, such as those implemented by middle frontal gyrus and ventromedial prefrontal cortex, project more ventromedially (Haber, Fudge, & McFarland, 2000). This organization has been replicated in humans using resting state functional connectivity (Marquand, Haak, & Beckmann, 2017). However, cortical projections that terminate along the lateral and medial walls of the caudate have been observed in nonhuman primates (Selemon & Goldman-Rakic, 1985; Yeterian & Van Hoesen, 1978). Indeed, recent work suggests the existence of a medial-lateral organizational scheme in the caudate that is not evident in the putamen (O’Rawe, Ide, & Leung, 2019). It thus seems necessary to further examine the organization of the caudate separately.

Methods that allow the continuous sampling of data across space not only allow using the space as a continuous factor, but afford simultaneous estimates of various organizational schemes within the same individual. Resting-state fMRI offers this capability and affords non-invasive measures of functional organization. While fMRI data have been utilized to reveal broadly the anterior-posterior organization of the caudate (Jaspers, Balsters, Kassraian Fard, Mantini, & Wenderoth, 2017; Manza et al., 2015; Martino et al., 2008), these previous studies, however, have not taken advantage of the continuous nature of the sampling space of fMRI data. More modern approaches attempt to delineate continuous forms of organization using gradient techniques and/or by using space as a factor of analysis (Haak, Marquand, & Beckmann, 2017; Jarbo & Verstynen, 2015; Marquand et al., 2017; O’Rawe et al., 2019).

In this study, we use space as a factor to examine linearly dependent changes in caudate functional connectivity across its anatomical space using resting-state fMRI data. This will allow us to identify and characterize at least the three potential orthogonal organizations: medial-lateral (M-L), anterior-posterior (A-P), or dorsal-ventral (D-V). Based on the previous literature, we expected to find two gradients of functional organization across the caudate’s anatomical space, A-P and M-L. To examine individual variations and the potential implications of caudate organization, we further explored the effects of aging on the caudate’s spatial organization and its relationship with age-related reductions in flexible behavior (Wecker, Kramer, Hallam, & Delis, 2005). Because both goal-directed behavior and striatal dopamine decline with age (Fearnley & Lees, 1991; Volkow et al., 1998), it has been hypothesized that this striatal dopamine loss is at least in part related to the age related decline in flexible behavior. Reduction in both striatal and midbrain dopamine was found to be correlated with a reduction in cognitive flexibility in heathy aging (Berry et al., 2016; Bäckman et al., 2000; Dreher, Meyer-Lindenberg, Kohn, & Berman, 2008)(Bäckman et al., 2000). As recent work has demonstrated that dopamine signals entering the striatum are spatially biased and sweep along the M-L direction (Hamid, Frank, & Moore, 2019), it is expected that we will find evidence of an aging-related reduction in caudate spatial organization (particularly along the M-L axis) due to a loss of this spatially biased signal. We further hypothesized that this decrease in spatial organization would be related to age related decreases in flexible behavior.

## Methods

### Imaging data and image acquisition parameters

For the primary caudate topography analyses, we utilized the Cambridge Buckner sample of 198 subjects (123 female from the 1000 Connectomes Project (Biswal et al., 2010) (http://fcon_1000.projects.nitrc.org), ages 18-30 (M = 21.03, SD = 2.31). For replication and age-related analyses, we utilized the 207 subjects (87 female) in the Rockland dataset (http://fcon_1000.projects.nitrc.org/indi/pro/nki.html), ages 4-85 (M = 35.00, SD = 20.00) and the first (in order of release) 377 human subjects (238 female) of the Rockland Enhanced dataset, ages 8-85 (M = 42.11, SD = 20.34). All datasets from the Cambridge Buckner database passed the screening for motion other image artifacts (with at least 2/3 usable data per subject). For the Rockland dataset 18 subjects were excluded, leaving 189 subjects (78 female) remaining, ages 4-85 (M = 35.70, SD = 19.89). For the Rockland enhanced dataset, 78 subjects were excluded, leaving 299 subjects (194 female), ages 8-85 (M = 40.96, SD = 20.33).

The Cambridge Buckner data (Siemens 3T Trim Trio): T1-weighted images were collected with MPRAGE with the following image parameters: slices = 192, matrix size = 144 × 192, voxel resolution = 1.20 × 1.00 × 1.33 mm^3^. Resting state T2*-weighted images were acquired using EPI with the following parameters: 47 interleaved axial slices, TR = 3000 ms, voxel resolution = 3.0 x 3.0 × 3.0 mm^3^ (119 volumes).

The NKI/Rockland data (Siemens 3T Trim Trio): T1-weighted images were collected using MPRAGE with the following parameters: slices = 192, matrix size = 256 × 256, resolution = 1.00 × 1.0 × 1.0 mm^3^. Resting state T2*-weighted images were acquired using EPI with the following parameters: 38 interleaved axial slices, slice gap = 0.33 mm, TR = 2500 ms, TE = 30 ms, Flip Angle = 80 deg, voxel resolution = 3.0 × 3.0 × 3.0 mm^3^ (260 volumes).

Enhanced Rockland (Siemens 3T Trim Trio): T1-weighted images were collected using MPRAGE with the following parameters: slices = 176, matrix size = 250 × 250, resolution = 1.00 × 1.0 × 1.0 mm^3^. Resting state fMRI data were acquired with the following parameters: Multi-band Acceleration Factor = 4, 64 interleaved axial slices, slice gap = 0 mm, TR = 1400 ms, TE = 30 ms, Flip Angle = 65 deg, voxel resolution = 2.0 × 2.0 × 2.0 mm^3^ (404 volumes in total).

### Image Preprocessing

Prior to analysis images were preprocessed utilizing SPM12 (http://www.fil.ion.ucl.ac.uk/spm/software/spm12/). Images were first corrected for slice timing, and then realigned to the middle volume according to a 6 parameter rigid body transformation. Structural images were coregistered with the mean functional image, segmented, and then normalized to the MNI template using both linear and nonlinear transformations. Functional images were then normalized utilizing the same parameters as the structural normalization.

For resting-state analysis, additional preprocessing steps were performed using either CONN (http://www.alfnie.com/software/conn) or custom Matlab scripts. A nuisance regression was constructed to regress out the following confounding variables: 6 motion parameters up to their second derivatives, scans with excessive motion, effects of session onset, modeled physiological signal generated through aCompCor (Behzadi, Restom, Liau, & Liu, 2007) of the white matter and CSF voxels, and a linear drift component. For the Cambridge Buckner and Rockland datasets, the residuals of this regression were filtered using a bandpass filter (.008 < *f* <.09), while for the Rockland Enhanced dataset this step was done simultaneously with regression (Hallquist, Hwang, & Luna, 2013). Finally, the residuals were despiked using a tangent squashing function.

### Spatially dependent functional connectivity pipeline

We named the pipeline Fine Relational Spatial Topography (FiRST). The goal of FiRST is to systematically explore spatial biases in functional connectivity of a region of interest. For instance, in this study, we asked whether or not certain areas of the brain are more strongly functionally coupled to the medial aspect of the caudate in comparison to the lateral aspect of the caudate. To do this, we constructed a multiple regression to predict the caudate’s voxel-wise connectivity with the rest of the brain using some Euclidean description of the space parameters for each of voxel in the caudate. If the strength of connectivity for a particular brain voxel changed linearly along a certain spatial dimension (M-L, A-P, or D-V) of the caudate, then the model would produce a large coefficient for that dimension for that brain voxel.

Figure 1 shows the schematic diagram for the FiRST pipeline. For each voxel within the caudate ROI, we calculated its functional connectivity with the rest of the voxels in the brain (via Pearson correlation of the timeseries). The correlation values of each voxel in the whole brain connectivity maps were vectorized and then transformed with Fisher’s Z transformation to reduce skewness due to bounded quantities (Fig. 1A). For each voxel in the brain, we fit a linear regression (standardized ordinary least squares) of that voxel’s Fisher’s Z values with the coordinates of each of the seed voxels (Fig. 1B).

**Figure 1:**
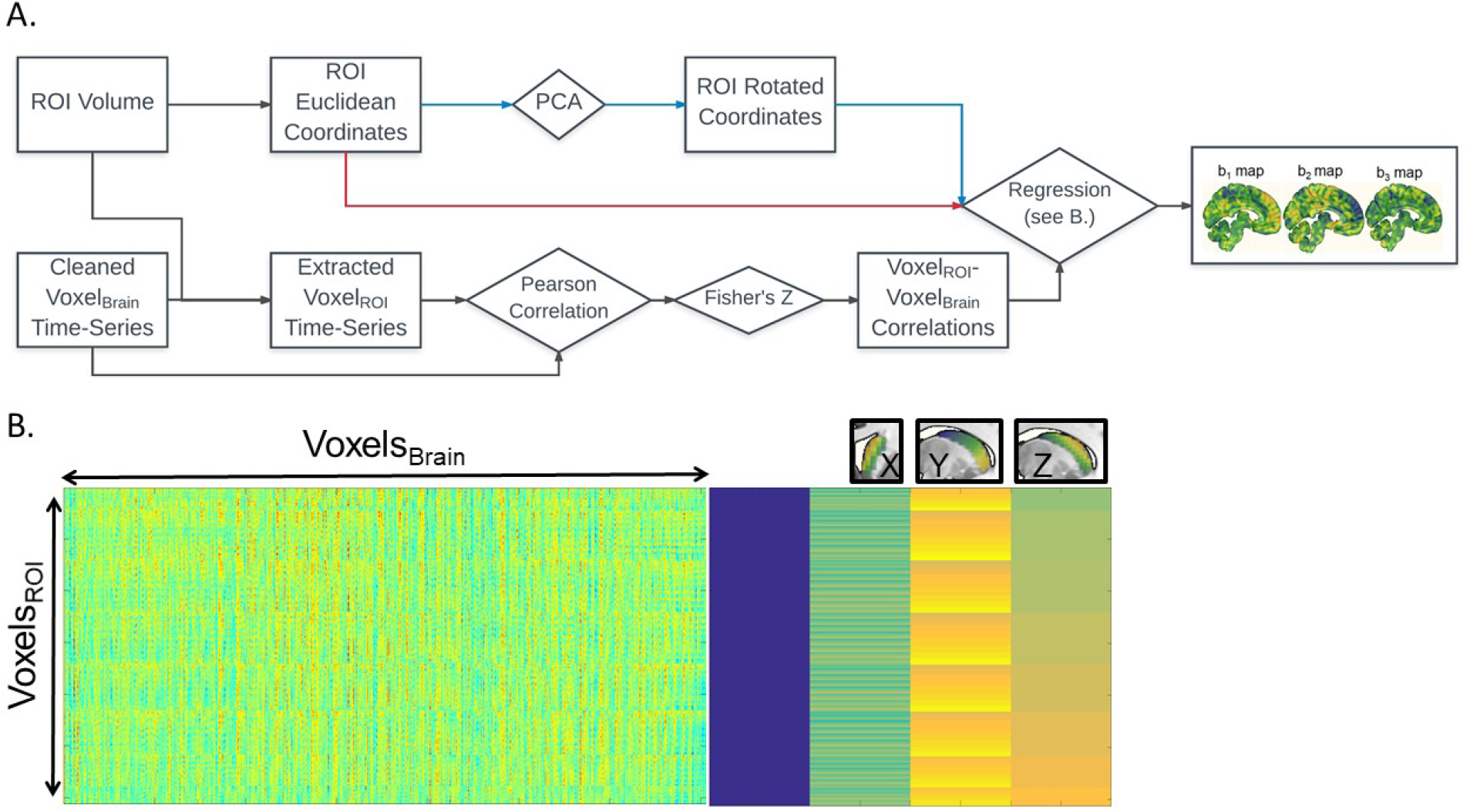
Fine Relational Spatial Topography (FiRST) approach. A. Analysis pipeline; squares denote measurements and diamonds denote operations. The voxel-wise time-series from the entire brain (Voxel_Brain_) are correlated with the voxel-wise time-series of the ROI (Voxel_ROI_). The resultant k × n matrix (where k = number of voxels in the ROI, and n = number of voxels in the brain) is then regressed against the coordinate space of the ROI, either in Euclidean coordinates or their corresponding rotated values derived from PCA (red or blue arrows in the flowchart). This regression produces 3 beta maps, denoting preference in connectivity along a particular directional gradient along the ROI (b_1_ for M-L, b_2_ for A-P, b_3_ for D-V). B. The visualization of the design matrix for the regression analysis. The rightmost side is equivalent to the conventional SPM design matrix, while the leftmost side is the input into the regression.

This process generates three β-weight maps representing change in connectivity strength (Fisher’s Z value) along each of the three Euclidean dimension (x, y, and z) of the ROI for each voxel in the brain (Fig. 1A right); that is, three spatial gradient connectivity maps (M-L, A-P, and D-V) for the caudate in each hemisphere. Note, prior to this regression step we orthogonalized caudate’s coordinate space utilizing Principal Component Analysis (PCA), a form of whitening. A rotation of any object without perfect symmetry within Euclidean space would induce a correlation between its coordinate space and a loss of spatial variance when performing regression. Orthogonalizing the space prior to regression can recover the spatial variance, though the dimensions may no longer follow traditional axes, depending on the extent of the rotation and the anatomical literature’s definitions. The FiRST pipeline has been released in the form of a general purpose toolbox and can be openly accessed via the NITRC (https://www.nitrc.org/projects/first/).

Second level t-tests were then conducted to generate group level gradient connectivity maps. We further calculated disjunction maps across the second level gradient maps in order to select voxels that demonstrated a strict bias across only one of the dimensions (e.g. a voxel has a significant (α = .001) negative beta in the medial-lateral axis but is no different than zero in either the anterior-posterior or dorsal-ventral axes). We further restricted the results to voxels that were above threshold in 2/3 of the datasets analyzed (“replication maps”).

### Age Regression Analysis

A whole brain regression analysis of each subject’s gradient β-weight maps was constructed for each orthogonal dimension. To test if the prototypical organization of cortical connectivity with the caudate degrades across age (i.e. gradient β-weights trending towards zero), we extracted age β-weights from the significant voxels on each side of the second level gradient disjunction map. If we are observing an overall degradation of organization, then the positive side will have a net negative relationship with age, while the negative side will have a net positive relationship with age.

### Estimations of Gradient Integrity

In order to more directly estimation the integrity of each gradient, we used a set of techniques that were subsequently developed based our prior gradient decomposition work in the striatum (O’Rawe et al., 2019). We previously found two predominant gradients of organization in the caudate, a M-L organization that resembles the othogonalized M-L dimensions here, and a dorsolateral to ventromedial diagonal gradient that most resembles the orthogonalized A-P axis here. From this prior work, we used the discovered gradients as gradient templates. First, to demonstrate the similarity of the results using the orthogonalized coordinate system and those using the prior templates, we examined the whole brain relationship of connectivity along each of these gradient templates. To do this, we ran a voxelwise regression (same as described above for the FiRST pipeline) with the templates as regressors instead of orthogonalized coordinates. The results are highly similar, as shown by the whole brain maps displayed in Supplemental Figure 1. Then, to construct an integrity measure, we fit each subject’s estimated gradients (estimated using nonmetric multidimensional scaling) to the two template gradients using a multivariate multiple regression:

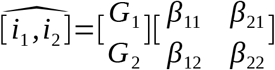

We then took the maximum absolute value beta for the fit of each gradient template as a measure the gradient integrity, with the intuition from previous work that each estimated space more often represents a single gradient.

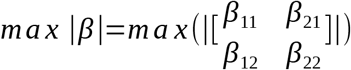

We also constructed an alternative metric without this assumption, the sum of the absolute value beta fit of each gradient template. This metric correlated heavily with the max metric and the results of all later tests were the same using either metric (only results from max metric shown).

### Testing the Relationship Between Caudate Gradient Integrity and Aging Related Changes in Executive Function

We used Partial-Least Squares (PLS) regression, a technique that derives latent variables with maximal covariance between sets of data, in the Rockland datasample in order to uncover age related trends in an executive function battery, the Delis–Kaplan Executive Function System (D-KEFS). The age matrix was comprised of a polynomial expansion of age, up to a 3^rd^ degree polynomial, while the D-KEFS matrix incorporated the 18 variables from the following sub-tests: trail-making, verbal fluency, design fluency, color-word interference (Stroop), sorting, twenty questions, word context, tower, and proverb tests. We produced 3 latent executive function factors, each of which we examined the relationship with the integrity of both the caudate M-L and diagonal gradients.

## Results

We used a spatial regression approach to demonstrate the dependence of the functional connectivity patterns of caudate on its own spatial geometry (see Fig. 1). For each voxel in the whole brain, this analysis generated an estimate of the slope of the functional connectivity along each of the three orthogonal axes of the caudate ROI. Using three datasets of a combined total 686 subjects (198, 189, and 299), we found reliable patterns of preferred functional connectivity across the geometry of the caudate (Fig. 2). As both left and right caudate analyses yielded similar results, the results and discussion are described only for the right caudate for simplicity.

**Figure 2:**
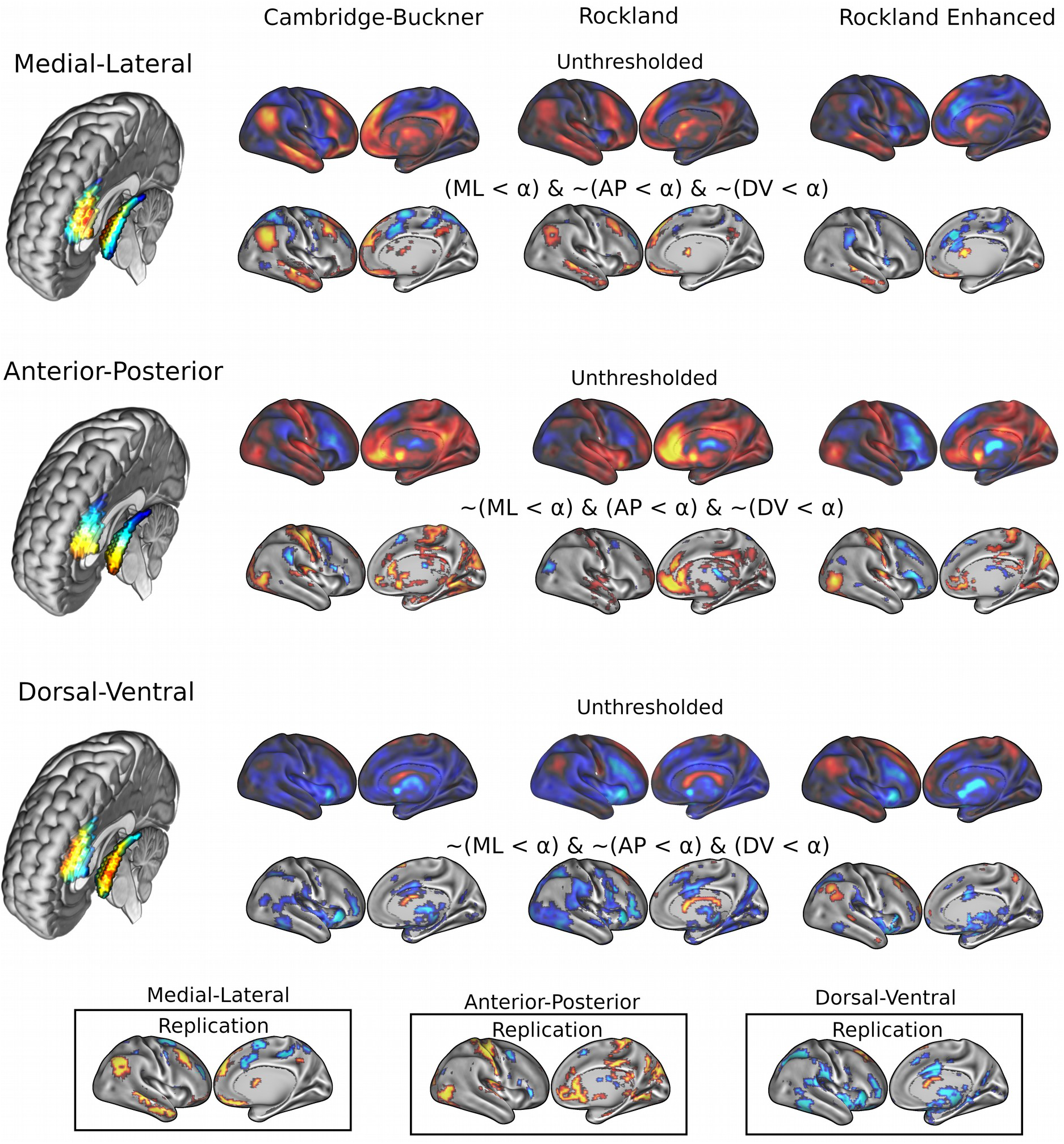
Results of FiRST analysis on caudate functional connectivity across 3 datasets: Cambridge-Buckner, Rockland, and Rockland Enhanced. For each datasample, we calculated the spatial betas uniquely related to each orthogonal dimension: M-L, A-P, and D-V. In the bottom panels, we calculated the replication maps across the three datasamples, where a voxel was maintained if it surpassed threshold in 2/3 datasamples.

### Gradient Connectivity along three main axes: M-L, A-P, and D-V

Figure 2 (top two rows) shows the whole-brain connectivity for the right caudate’s M-L gradient. The medial extent (warmer colors) of caudate demonstrated preferential connectivity with the dorsomedial and ventromedial prefrontal cortex (dmPFC and vmPFC), along with the posterior cinculate cortex/precuneus (PCC/Pcu), angular gyrus, and middle temporal gyrus; these regions are commonly referred to as part of the default mode network (DMN) (Greicius, Krasnow, Reiss, & Menon, 2003; Raichle et al., 2001). Frontoparietal areas including the inferior frontal gyrus/frontopolar cortex, posterior dorsal portion of the middle frontal gyrus and cerebellum also demonstrated preferential medial caudate connectivity. The lateral extent (cooler colors) of the caudate demonstrated preferential connectivity for the middle frontal gyrus (more anterior), superior frontal sulcus/dorsal aspect of FEF, frontal operculum, supplemental motor area (SMA) extending to pre-SMA, premotor, intraparietal sulcus, and superior parietal lobule; these regions are commonly referred to as part of the dorsal and ventral attention networks (Fox, Corbetta, Snyder, Vincent, & Raichle, 2006). Lateral connectivity preference was also found with regions of the cerebellum and putamen. As these maps revealed network-level organization, we conducted quantification of the M-L gradient connectivity at the network level using a 7 network partition from the literature (Yeo et al., 2011). Specifically, dorsal attention [Cambridge Buckner: t(197) = −4.60, p < .001; Rockland: t(188) = −1.39, p = .17; Rockland Enhanced: t(298) = −2.66, p < .01] and ventral attention [t(197) = −5.66, p < .001; t(188) = −3.33, p = .001; t(298) = −8.93, p < .001] networks demonstrated significant connectivity preference with the lateral caudate while limbic [t(197) = 3.99, p < . 001; t(188) = 2.45, p < .05; t(298) = 4.21, p < .001] and default mode network [t(197) = 10.52, p < . 001; t(188) = 5.60, p < .001; t(298) = 2.71, p < .01] demonstrated significant connectivity preference with the medial caudate.

A different pattern of functional connectivity gradient was observed for the caudate’s A-P axis. The anterior extent of caudate showed preferential connectivity with areas of sensorimotor, auditory, and visual cortices, along with pulvinar, posterior insular, precuneus, hippocampus, vmPFC/anterior cingulate, and superior frontal sulcus. More posterior portions of caudate displayed preferential connectivity with the posterior middle frontal gyrus, preSMA, and cerebellum (Fig. 2 middle). Quantification at the network level is shown in Figure 3 (middle). Specifically, visual [t(197) = 4.43, p < . 001; t(188) = 0.77, p = .44; t(298) = 3.60, p < .001] and sensorimotor [t(197) = 5.93, p < . 001; t(188) = 4.71, p < .001; t(298) = 5.67, p < .001] demonstrate significant connectivity preference with the anterior caudate, while dorsal attention network shows weak connectivity preference with the posterior caudate [t(197) = −1.34, p = . 18; t(189) = −2.20, p < .05, t(298) = −0.84, p = .40].

**Figure 3:**
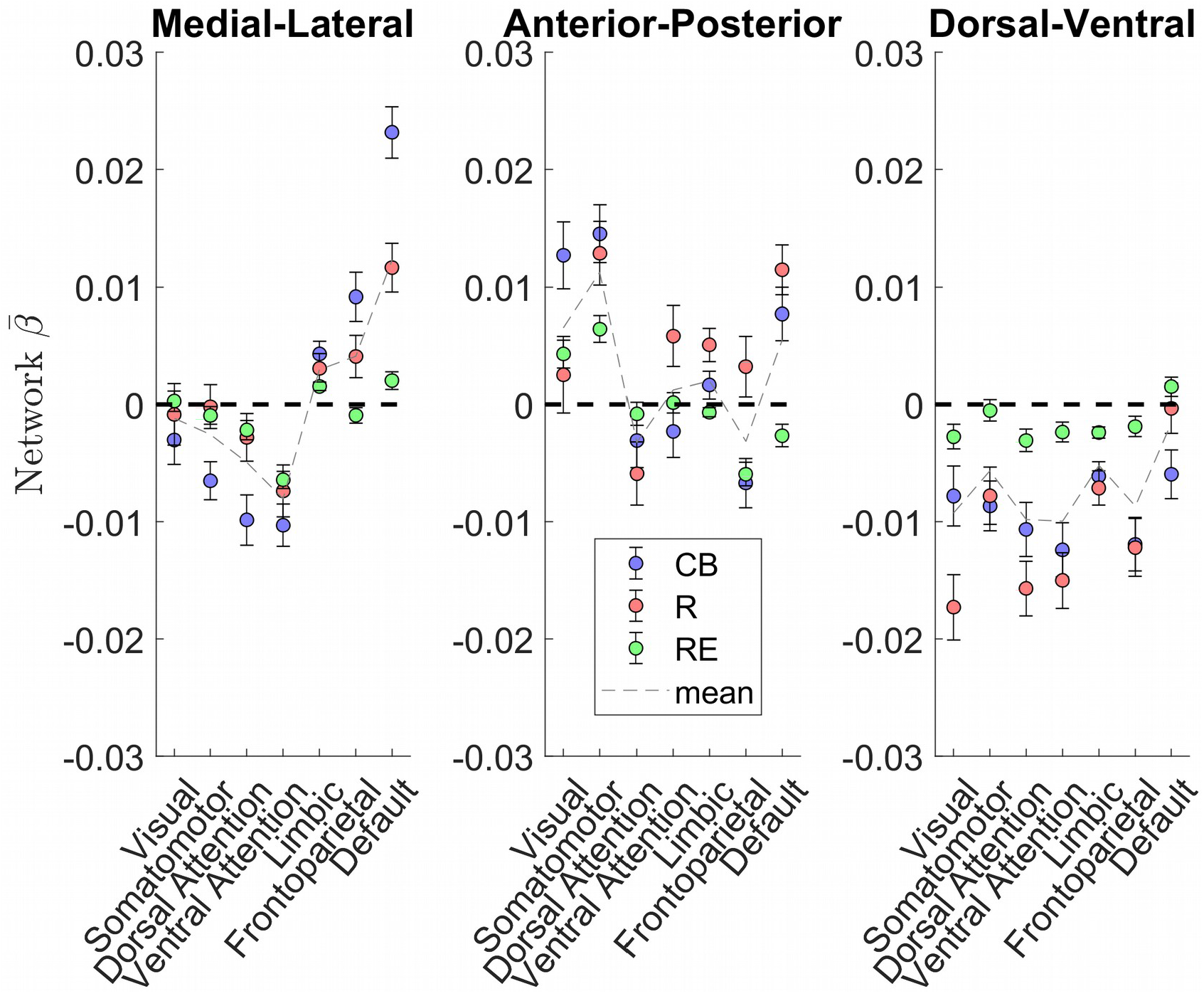
Network Estimation of caudate connectivity along the M-L, A-P, and D-V dimensions. The medial-lateral axis shows a clear distinction between default mode network preferring to connect along the medial aspect of the caudate, while the dorsal and ventral attention networks preferred to connect along the lateral aspect of the caudate. Visual and somatomotor networks preferentially connect to the anterior aspect of the caudate.

Unlike the two other gradients, the D-V gradient did not show clear biologically plausible patterns. The more dorsal portions of caudate showed some connectivity preference with the superior frontal gyrus, but along with voxels mostly in the ventricles and white matter. The more ventral portions of the caudate appeared to show some connectivity preference with across the large spans of cortex (Fig. 2 bottom). Supporting the interpretation of this gradient simply segregating areas of meaningful BOLD signal and areas of noise BOLD signal, the network extractions demonstrated a ventral preference for every network in the brain (Fig. 3 right).

### Aging, caudate topography, and cognitive decline

Age-related changes in caudate volume and connectivity have been shown and implicated in age-related decline in flexible behavior (Berry et al., 2016; Manza et al., 2015; Verstynen et al., 2012). We examined the relationship between age and caudate functional organization using the Rockland and Rockland Enhanced samples as they have a wide age range (4-85 years). Fig. 4A and 4B shows age related effects across the three spatial gradients. Specifically, we noticed that the spatial distribution of aging beta-weights were inversely related to the gradient weights, suggesting an aging related degradation of caudate functional organization (both positive and negative weights trend towards zero with age). To evaulate this observation more concretely, we selected voxels based on each dataset’s disjunction map, producing average aging betas from voxels that have a positive or negative spatial gradient beta (Fig 4C and 4D). Universally, across datasets and spatial gradients, the aging beta average is in the opposite direction of the gradient beta (i.e. negative for medial bias and positive for lateral bias).

**Figure 4:**
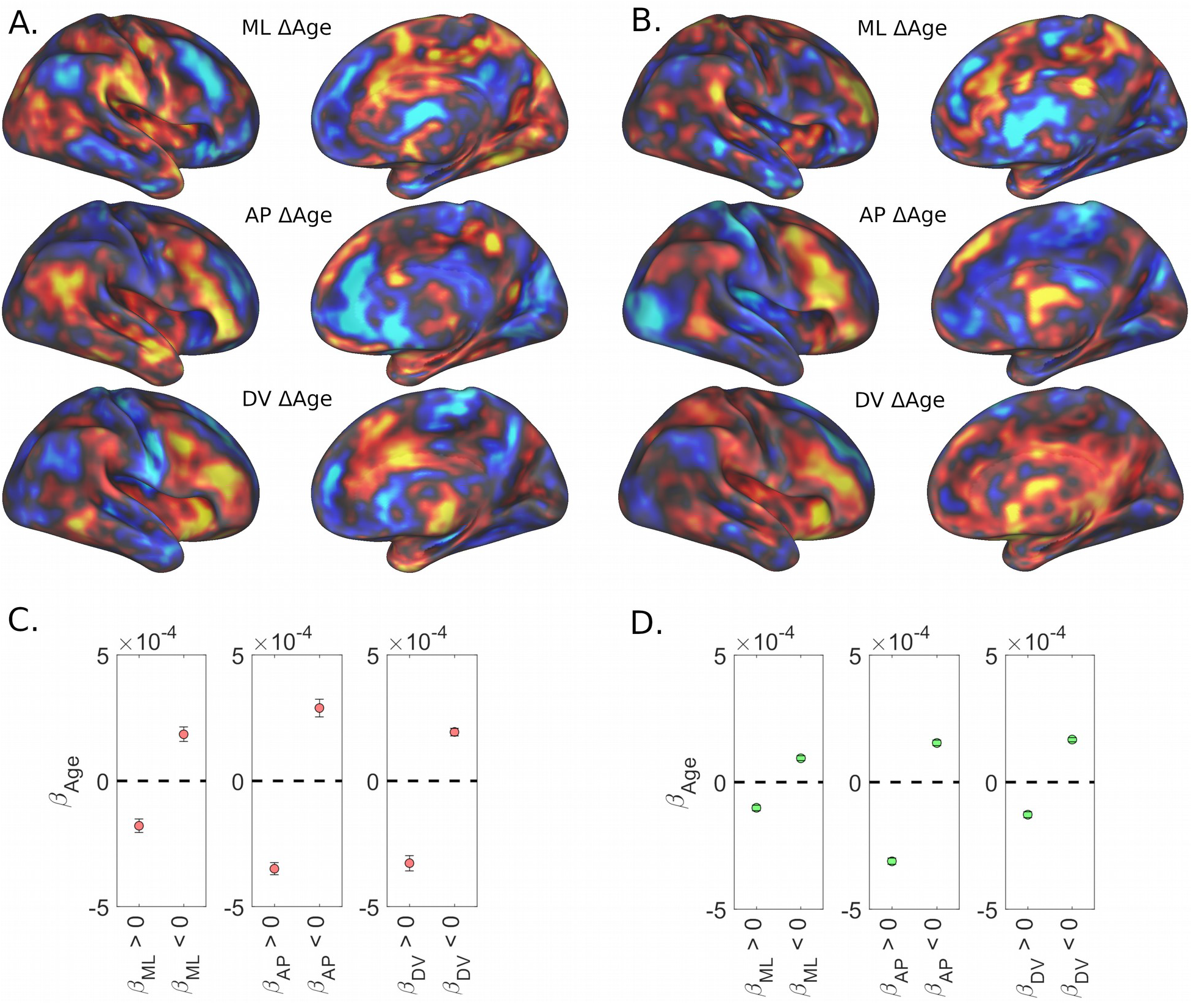
Aging Regression Results. **(A)** Aging regression results in the Rockland datasample presented unthresholded across the brain. **(B)** Aging regression results in the Rockland Enhanced datasample presented unthresholded across the brain **(C)** Average beta weight for age across significantly positive and significantly negative spatial weights in the Rockland datasample. **(D)** Average beta weight for age across significantly positive and significantly negative spatial weights in the Rockland Enhanced datasample.

Further, we used a new metric to compute each individual subject’s gradient integrity to relate to age, and examined to what extent that each gradient integrity contributes to age related changes in executive function. We used two gradient templates estimated in previous work (O’Rawe et al., 2019), medial-lateral (related to M-L axis here) and diagonal (related to A-P axis here) gradients, which showed similar cortical connectivity maps as the M-L and A-P gradients described above and shown in Figure 2 (Fig. S1). The fit of each subject’s estimated caudate gradients with the templates was used as a measure of gradient integrity (see Methods). We found a clear negative correlation with age for both gradients (ML: [r(187) = −0.34, p < .001; r(297) = −0.37, p < .001], Diagonal (or A-P): [r(187) = −0.20, p < .01; r(297) = −0.25, p < .001]) (Fig. 5).

**Figure 5:**
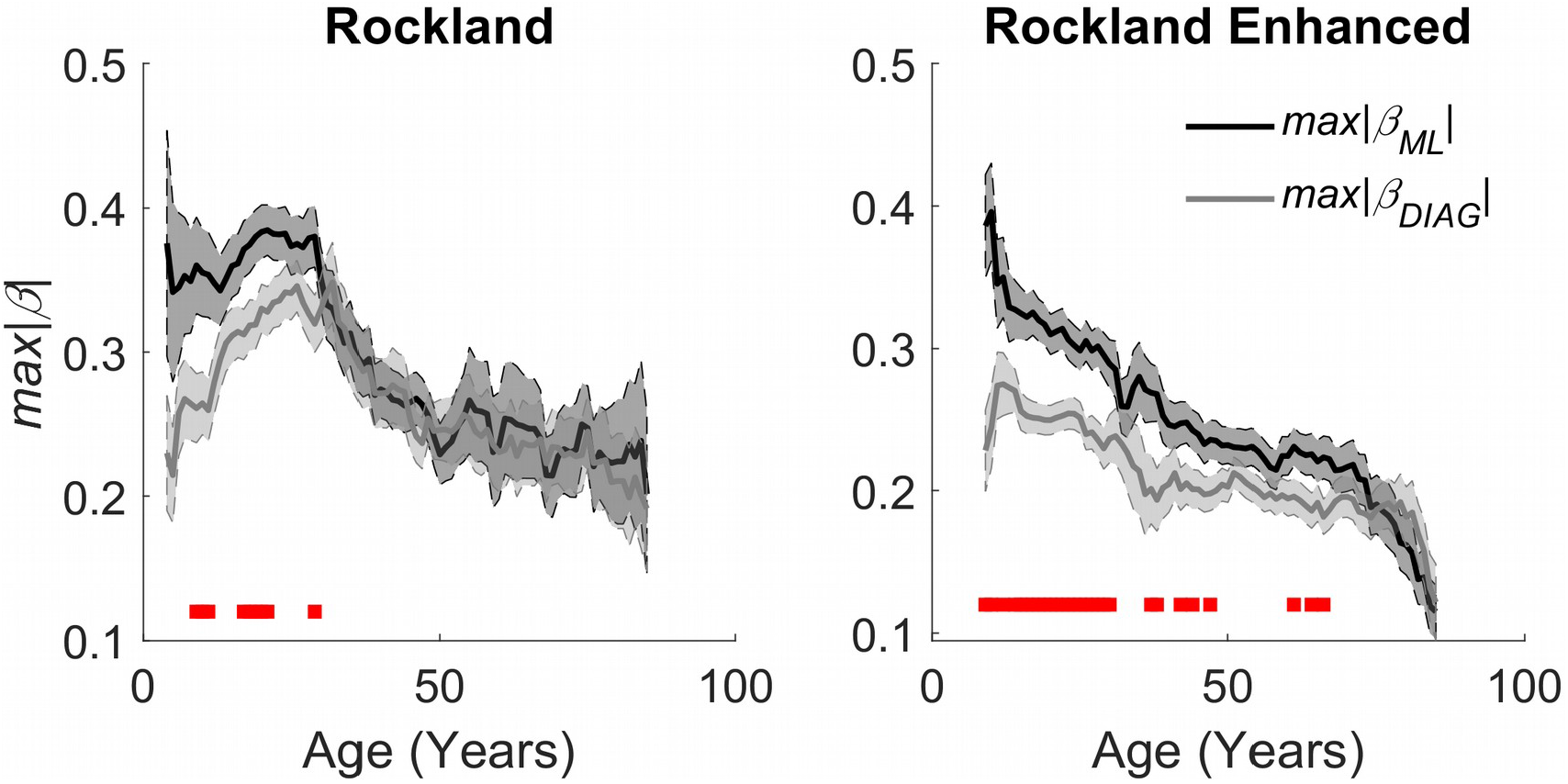
Aging effects on quantitative estimation of the integrity of caudate gradients (**Left)** Estimates across age groups using a sliding window mean of 14 years of the Rockland datasample. **(Right)** Replication of aging effects in the Rockland Enhanced database. Red bars at the bottom represent a significant difference between the two gradients with an alpha of .05, demonstrating that there’s a bias in organization across M-L early in age that evens out as the organization degrades with age.

To examine whether degradation of the caudate is related to aging related declines in executive function, we performed a partial least squares (PLS) regression on the Delis-Kaplan Executive Function System (D-KEFS) and a 3^rd^ order polynomial expansion of age in the Rockland sample and extracted 3 aging related latent executive function variables (Fig. 6A-C, Fig. S2 for loadings). We then examined whether these 3 latent executive function variables were related to estimates of either M-L or A-P gradient integrity. We found that the only relationship that surpassed FWE thresholds was the relationship between M-L gradient integrity and the PLS component 2 (r(187) = 0.24, p_FWE_ < .01) (Fig. 6D). This relationship was in part driven by the age related variance, as including age as a regressor in a linear model reduced the effect size to half its original size (β = 0.12, p = .12).

**Figure 6:**
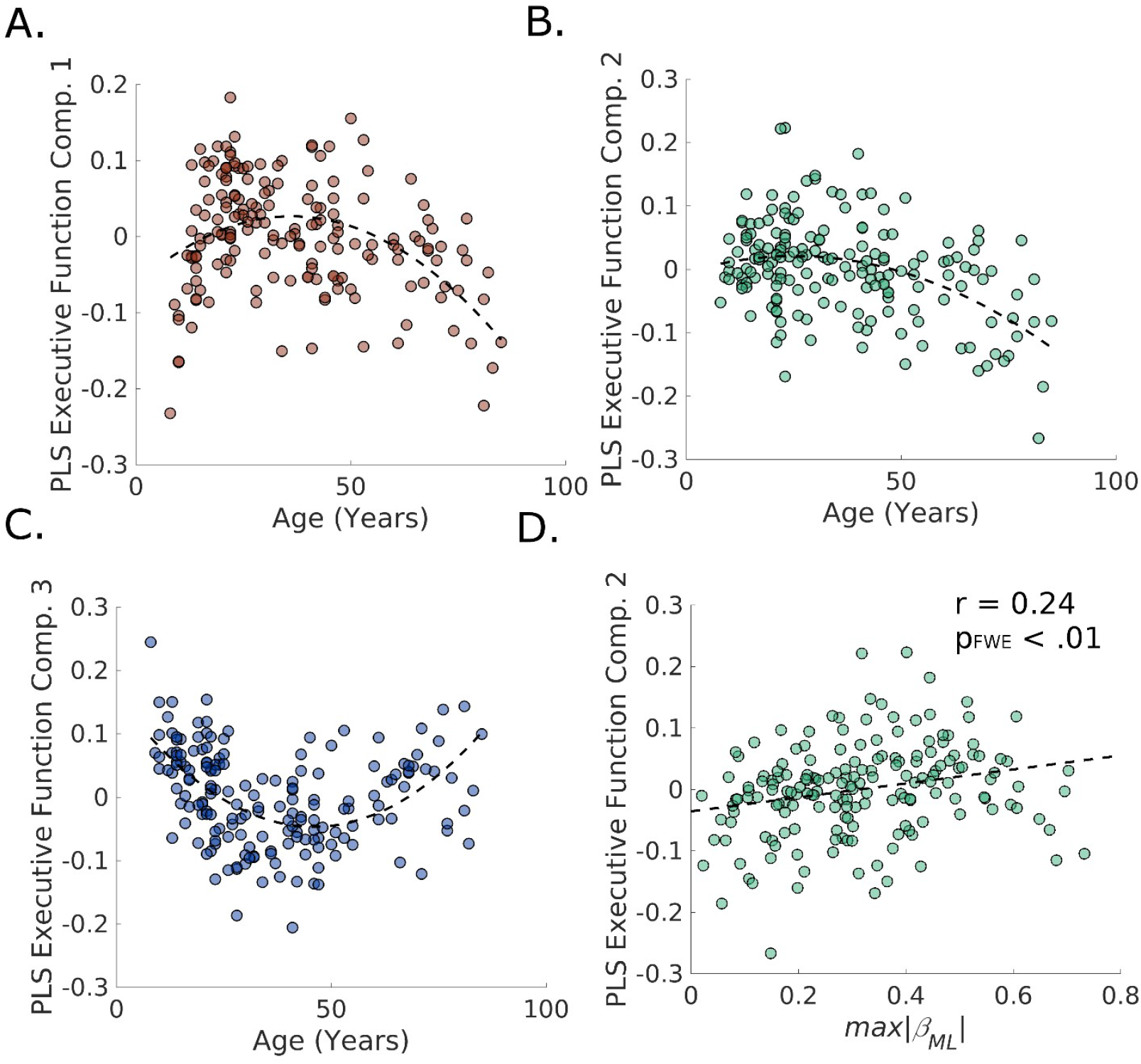
Results of PLS regression of age polynomial expansion (up to 3^rd^ order) on the D-KEFS inventory for the NKI/Rockland sample. **(A-C)** The three age-related latent executive function variables plotted across age. **(D)** The second age-related latent executive function variable significantly correlates with the integrity of the M-L gradient in the caudate.

## Discussion

While the caudate is posited as a central component of higher order neural circuits for supporting goal-directed behavior and action selection (Grahn et al., 2008; Mink, 1996), it is unclear what organizational features support such behavioral functions. Understanding the organizational layout of the cortico-caudate circuits may provide the basis for understanding. Utilizing the continuous space provided by resting-state fMRI data, we found a stable functional organization of the cortico-caudate topography along both the A-P and M-L axes (but not DV) of the caudate nucleus (Fig. 2). These topographic organizations, particularly the one along the M-L axis, appear to be organized by large scale networks (Fig. 3), as suggested in early anatomical tracing studies (Parthasarathy et al., 1992; Selemon & Goldman-Rakic, 1985; Yeterian & Van Hoesen, 1978). Furthermore, these caudate functional organizational principles seem to break down across age, potentially meaningful to age related changes in flexible behavior (Fig. 4,5,6).

Our findings highlight the potential importance of the M-L functional topography of the caudate nucleus in the human brain. Our findings revealed that the medial caudate seems to be differentially connected to default mode and limbic networks, while the lateral caudate seems to be differentially connected to the dorsal and ventral attention networks. These findings are consistent with the anatomical tracing literature in macaques, which showed that the medial caudate receives projections from orbitofrontal cortex, vmPFC, dmPFC, and superior temporal gyrus (STG) (Ferry et al., 2000; Selemon & Goldman-Rakic, 1985; Yeterian & Pandya, 1991; Yeterian & Van Hoesen, 1978), while the lateral caudate receives projections from intraparietal sulcus (IPS), inferior parietal lobule (IPL), frontal eye fields (FEF), and supplementary eye fields (SEF) (Parthasarathy et al., 1992; Selemon & Goldman-Rakic, 1985). It has been, however, unclear what exact functions, if any, this organization provides until recently. Morris and colleagues have recently found that the caudate constrains the effect of prediction error on causal inference by the observed covariances in the environment, via interactions with parietal lobe and medial prefrontal cortex (Morris, Dezfouli, Griffiths, Pelley, & Balleine, 2017). Our findings suggest that the medial-lateral organization may provide the substrate for the integration of prediction error and observed environmental covariances.

Alternately, the M-L associated network segregation may delineate processes depending on internal and external information, as previous literature has implicated default mode/limbic networks in controlling internally guided decisions and implicated dorsal/ventral attention networks controlling externally guided decisions (Nakao, Ohira, & Northoff, 2012). The striatum organizing inputs of these networks along the medial-lateral axis may provide a substrate for shifting between these two strategies. Given a recent literature of systematic modeling of reinforcement learning within complex environments (Leong, Radulescu, Daniel, DeWoskin, & Niv, 2017; Niv et al., 2015), it is possible that the caudate is integrating and leveraging information from attentional deployment and from internally learned rules. Interestingly, shifts in attention during reinforcement learning in complex environments correlates to increased activity in the networks identified as more related to lateral caudate, and the maintenance of values during this process correlates with activity in networks more related to the medial caudate (Leong et al., 2017; Niv et al., 2015). Within these models, older adults show a deficit in reinforcement learning, and they potentially compensate for this via increased selective attention to fewer environmental features (Radulescu, Daniel, & Niv, 2016), which is supported by our data suggesting a reduction in M-L organization within the caudate across age. This interpretation also is consistent with the interpretation that medial-lateral sweeps of dopamine function as spatially specific credit assignment, reinforcing circuits recently leveraged for successful behavior. In other words, this medial-lateral dopamine sweep is overlaid ontop of existing spatially discrete circuits (Hamid, Frank, & Moore, 2019).

In sum, we have demonstrated both an A-P and M-L organization of the caudate functional connectivity to the rest of the brain in large samples of human subjects. Our findings revealed systems level of functional organization involving the caudate and that this organization degrades with age in correspondence with a reduction in flexible behavior. Future studies will be necessary to understand how much of the organization is intrinsic to the striatum, and how much of it is dependent on dopaminergic input.

