## Supplemental Materials for "Topographic functional interactions between caudate and cortex are organized by network and degrade with age"

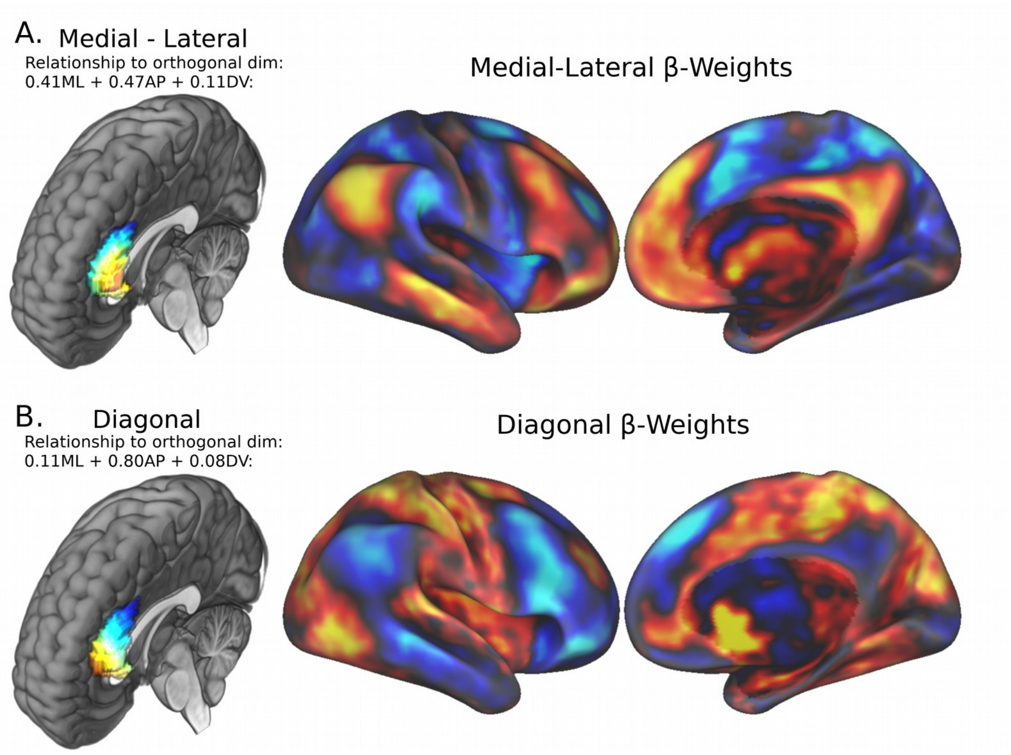


Figure S1: Caudate connectivity gradients derived using prior templates. Medial-lateral gradient **(A)** and Diagonal gradient **(B)**, as defined by previous empirically driven gradient discovery across the striatum (O'Rawe, Ide, and Leung, 2019), demonstrate similar caudate-cortical biases as the orthogonal M-L and A-P dimensions shown in Figure 2.

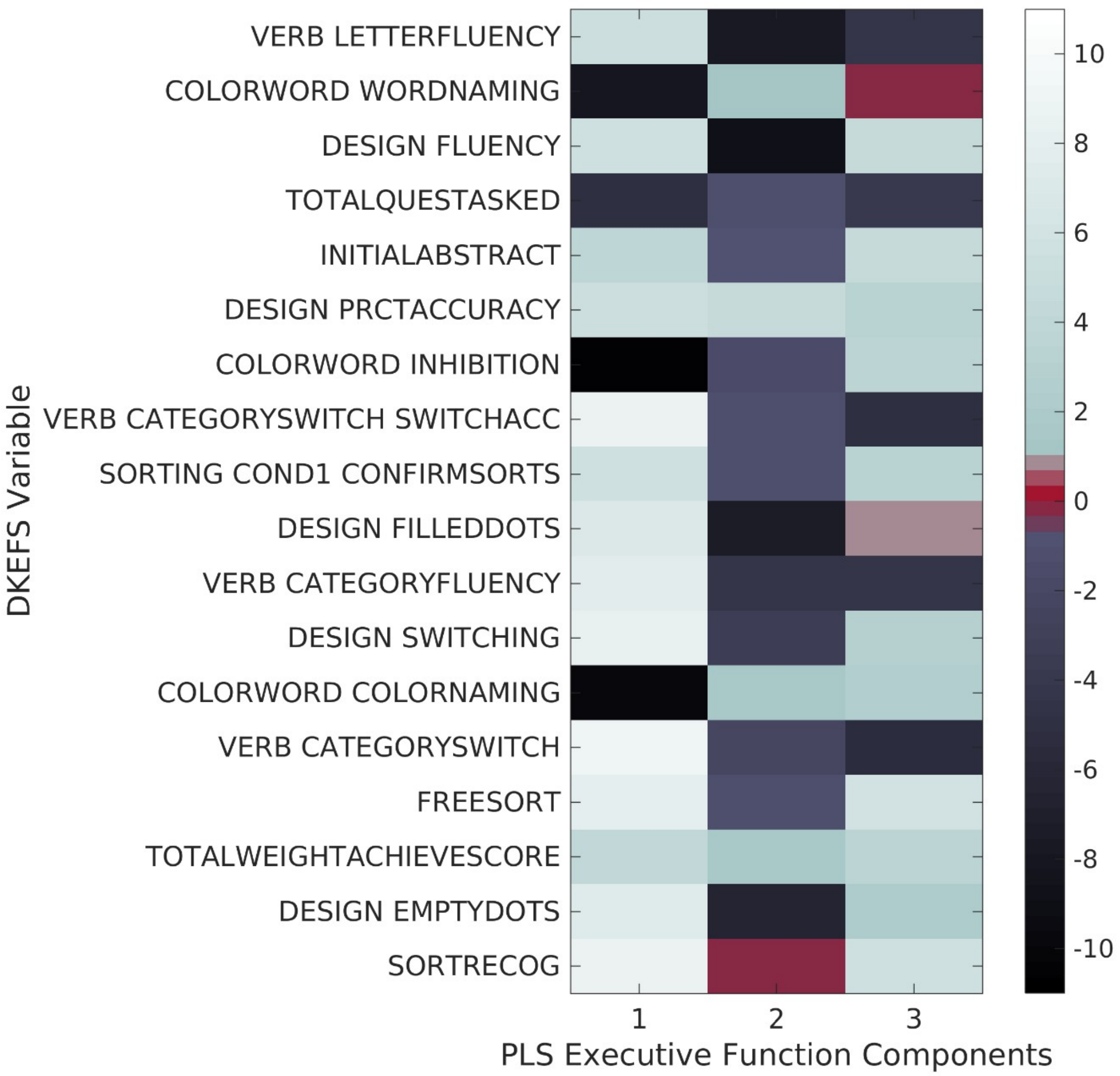


Figure S2: D-KEFS PLS loading matrix. Brighter colors denote larger positive loadings, darker colors denote larger negative loading. Note, positive PLS weights for Component 1 suggest a dependence on switching, composed of both verbal and visuospatial tasks, while the negative PLS weights for Component 3 seem more related to switching in the verbal domain.
